# Proteasome condensate formation is driven by multivalent interactions with shuttle factors and K48-linked ubiquitin chains

**DOI:** 10.1101/2023.06.25.546446

**Authors:** Kenrick A. Waite, Gabrielle Vontz, Stella Y. Lee, Jeroen Roelofs

**Author notes:** To whom correspondence should be addressed: Jeroen Roelofs, Department of Biochemistry and Molecular Biology, University of Kansas Medical Center, Kansas City, 3901 Rainbow Blvd., HLSIC 1077, Kansas, USA. equal contributions (Kenrick A. Waite, Gabrielle Vontz, Stella Y. Lee).

## Abstract

Stress conditions can cause the relocalization of proteasomes to condensates in yeast and mammalian cells. The interactions that facilitate the formation of proteasome condensates, however, are unclear. Here, we show that the formation of proteasome condensates in yeast depends on long K48-linked ubiquitin chains together with the proteasome shuttle factors Rad23 and Dsk2. These shuttle factors colocalize to these condensates. Strains deleted for the third shuttle factor gene, *DDI1*, show proteasome condensates in the absence of cellular stress, consistent with the accumulation of substrates with long K48-linked ubiquitin chains that accumulate in this mutant. We propose a model where the long K48-linked ubiquitin chains function as a scaffold for the ubiquitin binding domains of the shuttle factors and the proteasome, allowing for the multivalent interactions that further drive condensate formation. Indeed, we determined different intrinsic ubiquitin receptors of the proteasome (Rpn1, Rpn10, and Rpn13) are critical under different condensate inducing conditions. In all, our data support a model where the cellular accumulation of substrates with long ubiquitin chains, potentially due to reduced cellular energy, allows for proteasome condensate formation. This suggests that proteasome condensates are not simply for proteasome storage, but function to sequester soluble ubiquitinated substrates together with inactive proteasomes.

**Significance:** Stress conditions can cause the relocalization of proteasomes to condensates in yeast as well as mammalian cells. Our work shows that the formation of proteasome condensates in yeast depends on long K48-linked ubiquitin chains, the proteasome binding shuttle factors Rad23 and Dsk2 and proteasome intrinsic ubiquitin receptors. Here, different receptors are critical for different condensate inducers. These results indicate distinct condensates can form with specific functionality. Our identification of key factors involved in the process is crucial for understanding the function of proteasome relocalization to condensates. We propose that cellular accumulation of substrates with long ubiquitin chains results in the formation of condensates comprising those ubiquitinated substrates, proteasomes, and proteasome shuttle factors, where the ubiquitin chains serve as the scaffold for condensate formation.

## Introduction

Protein degradation mediated by the proteasome plays important roles in many cellular processes. Proteasomes have traditionally been considered as ready-to-go machines that are available whenever ubiquitinated substrates present themselves. They are also transcriptionally upregulated under conditions of proteolytic stress. However, more recently this view has been challenged because proteasome activity can also be modulated by post-translational modifications (1). Furthermore, superfluous or defective proteasomes can be targeted for autophagic degradation (2-5). Finally, proteasome localization is regulated in response to cellular changes, reflecting a localized need, e.g., in ER-associated degradation or surrounding aggregates. Interestingly, in yeast, proteasomes have been shown to be exported from the nucleus and to relocalize into distinct biomolecular condensates known as proteasome storage granules (PSGs) upon glucose starvation (6-8). Consistent with the liquid-liquid phase separation (LLPS) nature of these condensates, PSGs have liquid-like behavior and rapidly dissolve (within 10 minutes) upon re-addition of glucose to cells. While glucose utilization by fermenting single-celled organisms, like yeast, is a rather specialized response, recent work has shown that osmotic stress or amino acid starvation can induce proteasome LLPS in mammalian cells (9-11), suggesting there is a general cellular benefit to depositing proteasomes into condensates. Currently, it is unclear if the fundamental mechanism used by yeast and mammals to induce LLPS of proteasomes is conserved.

The yeast PSGs were proposed to be storage granules for proteasomes, as proteasomes rapidly returned to cell nuclei from cytoplasmic condensates upon glucose replenishment. Another model suggested proteasomes are sequestered into PSGs to protect them from incorporation into autophagosomes and subsequent degradation (8, 12). Here, however, it remains unclear why nuclear proteasomes would be prompted to leave the nucleus, as their nuclear localization also protects them from autophagy (2, 4). Regardless, both models propose that proteasomes are dissociated into regulatory particles (RP) and core particles (CP) before translocation to PSGs, together with monoubiquitin and the deubiquitinating enzyme Ubp6 (6-8, 12-14). However, these models do not provide a rationale for the multivalent forces required for the formation of proteasome condensates (15). Interestingly, in the mammalian system, proteasome condensates appear to actively degrade specific substrates (9, 11), providing an alternative function for proteasome relocalization into condensates. Here, formation of these condensates depended on one specific proteasome shuttle factor, RAD23B.

Proteasomes can bind ubiquitinated substrates either directly via three intrinsic receptors that have affinity for polyubiquitin chains (16), or indirectly using extrinsic receptors that can bind both ubiquitinated substrates and the proteasome (17, 18). The extrinsic receptors are known as shuttle factors and consist of three members in yeast, Dsk2, Rad23, and Ddi1, each with additional orthologs in humans (Fig. 1A). The shuttle factors are characterized by the presence of a Ubiquitin-Like (UBL) domain that binds proteasomal ubiquitin receptors and at least one Ubiquitin-Associated (UBA) domain that interacts with ubiquitinated substrates. While shuttle factors exhibit redundancy (particularly Rad23 and Dsk2), instances of substrate selectivity for each factor have been reported (19-22). Phenotypic evidence suggests shuttle factors are involved in a variety of cellular processes including cell cycle progression, spindle pole body duplication, and DNA damage responses (20, 23-25). Recent work shows ∼90% of proteasomal substrates are delivered to proteasomes by Dsk2 or Rad23 suggesting shuttle factors are prominent contributors to overall ubiquitin proteasome system (UPS) function (22). Although generally considered substrate transporters (18, 26-28), alternative models suggest shuttle factors protect ubiquitin signal integrity or function in direct opposition of proteolysis (29, 30). The latter could be achieved by stabilizing substrates through obstruction of ubiquitin chain elongation or sequestration of substrates away from degradation machinery (31-33). Thus, how shuttle factors influence proteasome dynamics is an emerging and important question.

**Figure 1.**
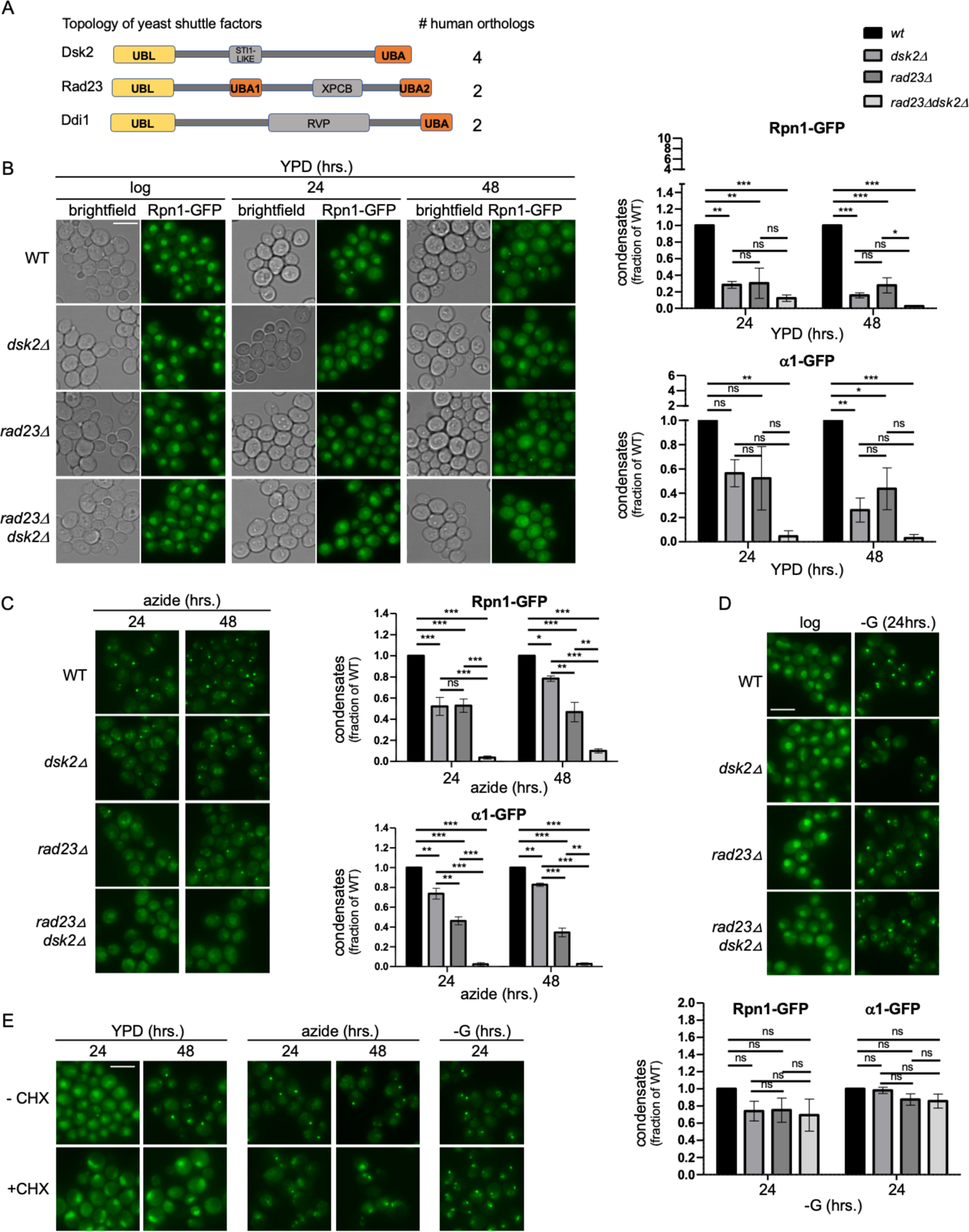
Rad23 and Dsk2 are important for the formation of proteasome condensates. **(A)** Topology of yeast shuttle factors and the number of human orthologues. UBL (ubiquitin like domain), UBA (ubiquitin associated domain), STI1 (stress inducible 1 domain), XPCB (XPC binding domain, RVP (retroviral protease domain). **(B)** Wildtype (WT), *dsk2*11, *rad23*11 and *rad23*11 *dsk2*11 yeast expressing Rpn1-GFP (microscopy images and quantification) or α1-GFP (quantification), tagged at their endogenous locus, were grown in yeast extract peptone dextrose (YPD) media and proteasome localization in live cells was visualized at indicated times using fluorescence microscopy. **(C)** Strains as in B were grown logarithmically, then treated with sodium azide for 24 or 48 hours before microscopic analysis of proteasome localization. **(D)** The indicated yeast strains were starved for glucose for 24 hours. Quantification shows fraction of cells that formed condensates relative to wild type cells. **(E)** Rpn1-GFP tagged cells were exposed to condensate inducing conditions in the presence or absence of cycloheximide. Proteasome localization in live cells was visualized using fluorescence microscopy at indicated time points. Quantification in (B), (C), and (D) was carried out using FIJI. The average of at least three biological repeats are shown and error bars represent standard deviation. 1-way ANOVA and Tukey’s multiple comparison test was used to determine significance (ns = not significant, * P < 0.05, ** P < 0.01, *** P < 0.001). The scale bars represent 5 μm.

In this report we show that shuttle factors play an important role in proteasome condensate formation in yeast, like they do in mammals. While these shuttle factors do not appear to be essential for condensate formation following glucose starvation, proteasome condensates induced by mitochondrial stress depend on Rad23 and Dsk2 for their formation. Like proteasomes, Rad23 and Dsk2 are enriched in these granules, suggesting these proteins have a direct role in proteasome condensate formation. Furthermore, we observed a dependance on K48-linked ubiquitin chains, but not K63-linked chains, in this condensate formation. Consistent with a role of ubiquitin chains as a scaffold in these condensates, mutations of intrinsic proteasome ubiquitin receptors disrupted condensate formation. In all, we propose a model where the accumulation of substrates with long K48-linked ubiquitin chains provides a prerequisite for proteasome condensate formation. Here, multivalent interactions between proteasomes, Rad23, Dsk2 and K48-linked ubiquitinated substrates trigger LLPS.

## Results

### Shuttle factors are required for proteasome condensate formation

While the shuttle factors are generally considered to deliver substrates to the proteasome, we were interested in testing if these factors play a role in relocalization of proteasomes. To explore this, we evaluated their role in proteaphagy as well as proteasome condensate formation. In mammalian cells, p62 has been identified as a proteaphagy adaptor linking proteasomes to autophagosome components (34). A p62 homolog does not exist in yeast, however, yeast shuttle factors and p62 share structural and functional similarities (namely, the ability to bind polyubiquitinated proteins and interact with proteasomes (35)). Furthermore, ubiquilins, the mammalian orthologs of yeast shuttle factor Dsk2, are important for general autophagy (36). To determine if shuttle factors were required for proteaphagy in yeast, we starved strains with GFP-tagged proteasome subunits, combined with deletions of the shuttle factors, for nitrogen or treated them with proteasome inhibitor (the latter only shows modest proteaphagy in our hands). We introduced a GFP-tag at the endogenous locus of RPN1, a regulatory particle subunit, or α1, a core particle subunit, such that every copy of this subunit is tagged and both subcomplexes can be monitored. Both subunits are essential, and the tagged subunit is efficiently incorporated into the proteasome (37). After 24 hrs. of nitrogen starvation or proteasome inhibitor treatment, the deletion mutants showed proteaphagy similar to wild type (WT), as was apparent from the accumulation of “free” GFP on immunoblots and a vacuolar GFP localization comparable to WT by fluorescence microscopy (Fig. S1). This indicates that, in yeast, shuttle factors are not required for efficient proteaphagy following nitrogen starvation or proteasome inhibitor treatment.

Next, we evaluated if the yeast shuttle factors were required for proteasome condensate formation. This was based on the observations that human ubiquilin-2 (UBQLN2), is capable of liquid-liquid phase separation (LLPS) and localizes to foci reminiscent of proteasome condensates (38), as well as the recently observed colocalization of human RAD23B in proteasome condensates (9, 11). The localization of GFP-tagged proteasomes was monitored in WT and shuttle factor mutant strains following growth in rich media (YPD) for an extended period (generally 48 hrs., here onwards referred to as prolonged growth). Consistent with previous reports, approximately 10% of WT cells exhibited cytosolic foci with proteasomes at 24 hrs. of growth in YPD (6). This percentage increased to ∼40% at 48 hrs. (Fig. 1B). The deletion of either *DSK2* or *RAD23* reduced the fraction of cells with condensates by ∼65-75% compared to WT cells at both time points (Fig. 1B). A deletion of both genes resulted in the complete absence of condensates, indicating a redundant role for Rad23 and Dsk2 in this process. Such a redundancy is not surprising and has been observed for other phenotypes as well (18, 22).

Some labs monitor proteasome condensates formation at stationary phase (quiescence), which involves growth for more than 5 days in YPD media (7). To determine if the absence of these shuttle factors caused a delay in condensate formation or the shuttle factors are essential for the process, we extended our prolonged growth in YPD up to 6 days. While the RAD23 and DSK2 single deletion strains showed some increase in the amount and intensity of condensates when comparing 3 and 6 days with 1 and 2 days, the lack of condensate formation in the *rad23*τ1 *dsk2*τ1 strain shows these two shuttle factors are required for this process and do not simply delay or decrease the efficiency of condensate formation (Fig. S2). In sum, our data indicate that both Dsk2 and Rad23 contribute to the localization of proteasomes in condensates under conditions of gradual nutrient depletion.

We recently reported that the treatment of cells with inhibitors of mitochondrial oxidative phosphorylation, like sodium azide, induced the formation of proteasome condensates (39). In this condition, the deletion of both RAD23 and DSK2 also resulted in an almost complete loss of condensate formation, indicating that the shuttle factors play a critical role in proteasome localization following mitochondrial inhibition/stress (Fig. 1C). Surprisingly, acute glucose starvation, another condition that results in PSG formation, did not show any measurable deficiencies in the formation of proteasome condensates in *rad23*τ1 *dsk2*τ1 cells (Fig. 1D). This contrasts with a recent study observing that *rad23*τ1 cells were defective in the formation of glucose starvation-induced condensates (11). This study, however, was mainly focused on mammalian cells and did not study the role of the yeast shuttle factors in detail. In addition to the differential requirement for shuttle factors in proteasome condensates formed by prolonged growth, sodium azide treatment or glucose starvation, we also observed a difference in the need for protein synthesis. Condensates that were dependent on Rad23 and Dsk2 for their formation, i.e. those induced by prolonged growth and azide treatment, also required protein synthesis, as the addition of the translation inhibitor cycloheximide disrupted their formation (Fig. 1E). In mammalian cells, cycloheximide treatment was shown to inhibit the formation of proteasome condensates known as starvation induced proteasome assemblies in the nucleus (SIPANs) because it prevented depletion of amino acid pools (11). However, this is unlikely to play a role here because yeast, unlike the mammalian cells, do not form proteasome condensates upon amino acid starvation (3). In all, we observed a striking difference in the requirements for condensates formed upon acute glucose starvation compared to other condensate inducing conditions, and Rad23 and Dsk2 are not a priori essential for the formation of proteasome condensates.

### Proteasomes, Rad23, and Dsk2 co-localize in condensates

Several factors have shown to be important for proteasome condensate formation, however they do not colocalize with proteasomes in the condensates, suggesting an indirect or regulatory role (7, 40). The requirement for Rad23 and Dsk2 in proteasome condensate formation and their ability to bind proteasomes suggests a direct role for these factors. To test this, we tagged the shuttle factors with mCherrry in our GFP-tagged proteasome strains. It should be noted that tagging Rad23 or Dsk2 reduced the number of condensates that formed, presumably because the tag compromises the function of these shuttle factors to some extent. Most condensates that were induced with azide or prolonged growth (both depending on Rad23 and Dsk2 for formation) showed colocalization of proteasomes with Rad23 or Dsk2 (Fig. 2). This is consistent with a direct role of the Dsk2 and Rad23 in the formation of these proteasomes. Interestingly, while the condensates that formed under glucose starvation did not depend on Rad23 or Dsk2, we observed that these shuttle factors still colocalized with proteasomes under those conditions (Fig. 2). In sum, the shuttle factors Rad23 and Dsk2 co-localize with proteasome condensates and are important for the formation of specific proteasome condensates.

**Figure 2.**
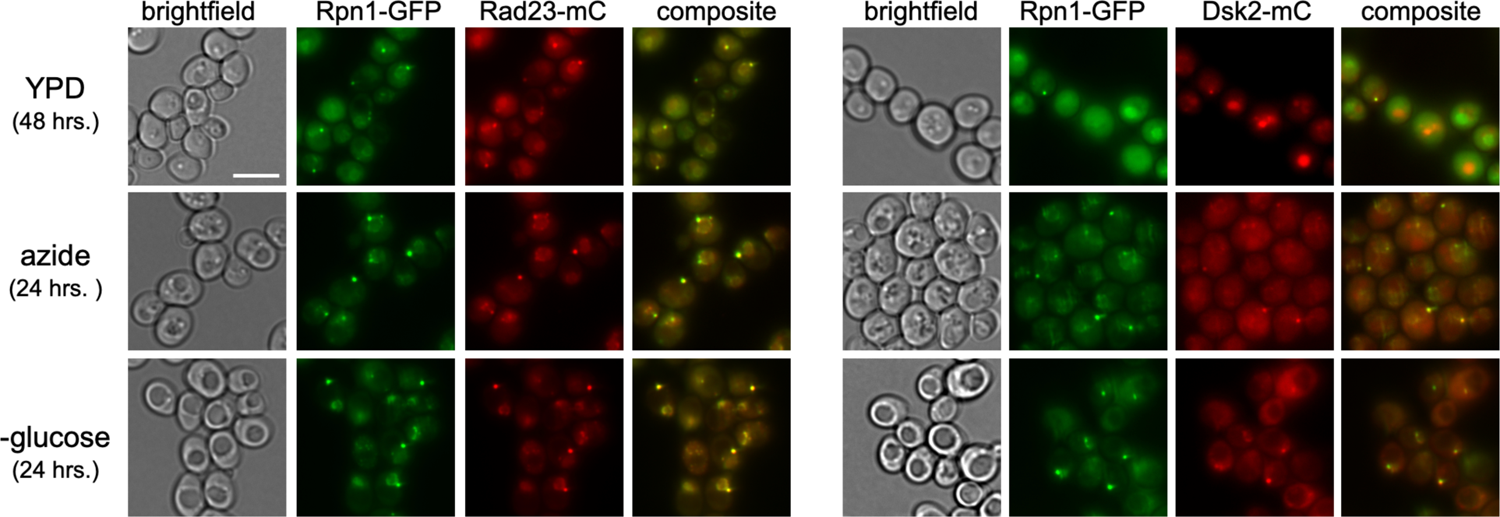
Rad23 and Dsk2 co-localize with proteasomes in condensates. Rad23-mCherry and Dsk2-mCherry were introduced into an Rpn1-GFP tagged strain to monitor their localization during prolonged growth, sodium azide treatment, and glucose starvation. The scale bars represent 5 μm.

### Ddi1 limits proteasome condensate formation

In addition to the shuttle factors Rad23 and Dsk2, there is a third shuttle factor, Ddi1. Similar to Rad23 and Dsk2, Ddi1 contains UBL and UBA domains for proteasome and substrate interactions, respectively. Like Rad23 and Dsk2, we did not observe a role for Ddi1 in proteaphagy (Figure S1). However, unlike Rad23 and Dsk2, we did not observe a strong dependence on Ddi1 for condensate formation during prolonged growth or sodium azide treatment and none of the shuttle factors was required for glucose starvation induced condensates (Fig. 3A, B). Instead, a deletion of *DDI1* substantially increased the fraction of cells with GFP-positive foci during prolonged growth. Furthermore, Ddi1 did not consistently co-localize with proteasome foci, although some co-localization was observed (Fig. 3C). Interestingly, we observed some Ddi1 foci where proteasomes were absent. The nature of these foci was not further investigated in this study.

**Figure 3.**
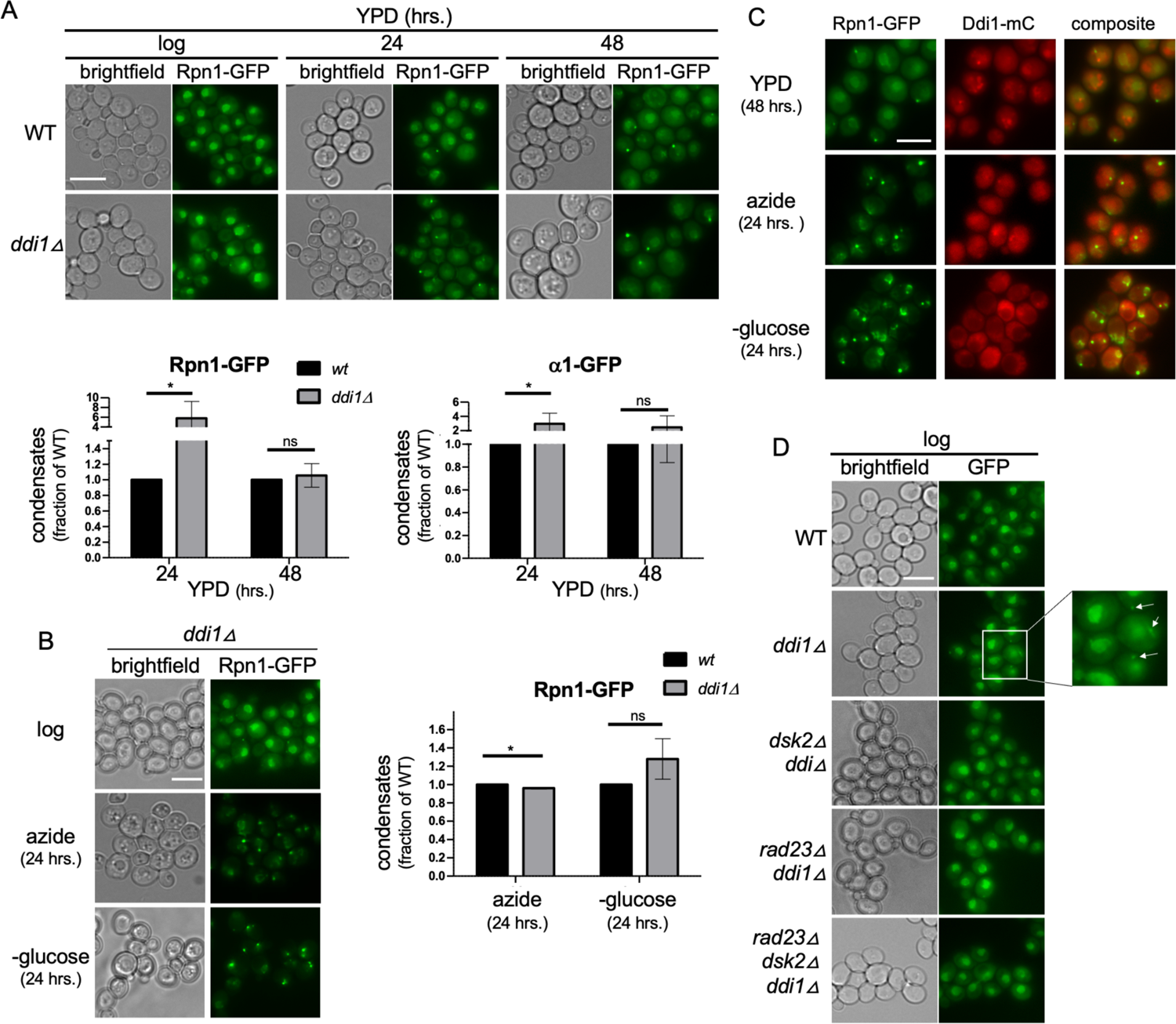
Ddi1 counteracts proteasome condensate formation. **(A)** Wildtype and *ddi111* cells expressing Rpn1-GFP were grown in YPD media and proteasome localization was monitored at the indicated times. Quantifications were carried out as in Fig. 1 and error bars represent standard deviation. T-tests were used to determine significance. **(B)** *ddi1 11* cells were monitored for a role of Ddi1 in proteasome condensate formation following sodium azide treatment or glucose starvation. Quantifications show this mutant relative to the wildtype strain shown in Fig. 1C and D. Standard deviation is presented, and T-tests were used to determine significance **(C)** Ddi1 was tagged with mCherry in an Rpn1-GFP tagged strain and its localization monitored at the indicated times and following sodium azide or glucose starvation for 24 hours. **(D)** Indicated strains, all Rpn1-GFP tagged, were grown logarithmically prior to fluorescent imaging. ns = not significant, * P < 0.05. The scale bars represent 5 μm.

While Ddi1 was not required for, nor consistently localized with, proteasome condensates, we did observe a striking phenotype in the DDI1 knockout cells growing logarithmically. That is, without any inducer or stressor, *ddi1Δ* cells formed proteasome condensates in log phase (Fig. 3D). These condensates were weaker in fluorescence intensity compared to the condensates observed with prolonged YPD growth and were still dependent on Rad23 and Dsk2 for their formation as they were absent in the *rad2311 dsk211 ddi111* cells (Fig. 3D). Thus, Ddi1 prevents the formation of cytosolic proteasome condensates in logarithmically growing yeast.

### Long K48-ubiquitin chains trigger proteasome condensate formation

Both human and yeast Ddi1 contain a viral protease domain that was recently shown to cleave substrates tagged with long polyubiquitin chains (41). This function of Ddi1potentially reflects a role in the cleavage and degradation of substrates that are difficult to degrade, e.g., due to the lack of an initiation site for proteasome engagement. In all, the presence of Ddi1 is critical to prevent the accumulation of substrates with long ubiquitin chains during logarithmic growth. Our observation that *ddi1*11 strains show proteasome condensates in logarithmically growing cells led us to hypothesize that the lack of Ddi1 protease activity, and consequently the accumulation of proteasome substrates in this mutant, trigger proteasome condensate formation. To test this, we overexpressed either wildtype Ddi1 or Ddi1^D220N^, a protease-dead mutant, in a DDI1 deletion background and monitored condensate formation. While expression of WT Ddi1 rescued the phenotype of proteasome condensates in logarithmically growing cells, expression of the protease-dead mutant did not (Fig. 4A). This shows that the protease activity of Ddi1 is required to prevent proteasome condensate formation during logarithmic growth. Thus, in *ddi1*11 cells condensate formation is not due to the loss of a competitor amongst the shuttle factors for proteasome binding, but instead likely reflects a role for the accumulation of proteasome substrates with long ubiquitin chains in condensate formation.

**Figure 4.**
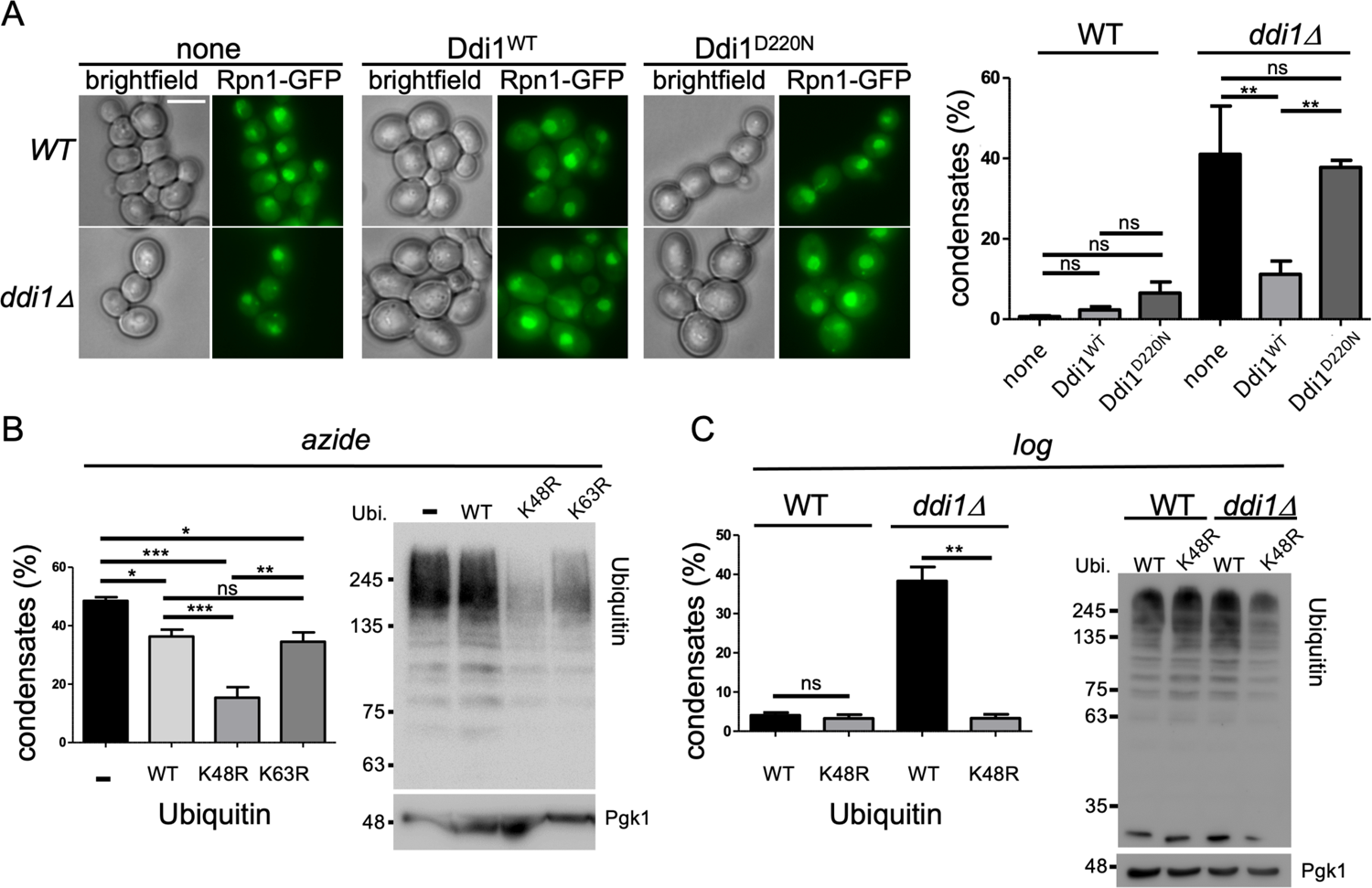
Long K48-linked ubiquitin chains are a critical component of proteasome condensates. **(A)** Logarithmically growing WT and *ddi1*11 yeast containing a plasmid without DDI open reading frame (none), expressing Ddi1 (Ddi1^WT^), or a protease dead Ddi1 (Ddi1^D220N^), were monitored by fluorescence microscopy for proteasome localization. Quantification shows the percentage of cells that formed proteasome condensates. 1-way ANOVA and Tukey’s multiple comparison test used to determine significance. **(B)** Wildtype cells expressing Rpn1-mCherry (-) were compared with strains overexpressing ubiquitin (WT), ubiquitin with lysine 48 mutated to arginine (K48R), and ubiquitin with lysine 63 mutated to arginine (K63R) following treatment with sodium azide. Ubiquitin variants were introduced at the URA-TIM9 locus to avoid issues with plasmid loss and expression was driven by a GPD promoter. The average of four independent experiments is shown with standard error of the mean. 1-way ANOVA and Tukey’s multiple comparison test was used to determine significance. Samples were collected after azide treatment and cell lysates were immunoblotted for ubiquitin and Pgk1. **(C)** Rpn1-GFP WT and *ddi1*11 yeast expressing ubiquitin (WT) or K48R ubiquitin were grown logarithmically and monitored for proteasome condensate presence (left). The average of four independent experiments is presented with standard error of the mean. Paired t-test was used to determine statistical significance. Cells were also collected lysed, followed by immunoblotting for ubiquitin and loading control Pgk1 (right panel). ns = not significant, * P < 0.05, ** P < 0.01, *** P < 0.001. The scale bar represents 5 μm.

Condensates generally rely on numerous low-affinity multivalent interactions and a scaffold to trigger their formation (15, 42). Based on the observed condensates in the *ddi1*11 strain, we reasoned that it is likely that the substrate-attached polyubiquitin chains form a scaffold for proteasome condensates to nucleate. We hypothesized that to function as a scaffold, a critical length needs to be reached to allow the ubiquitin chain to interact with the UBA domains of Rad23 or Dsk2 and/or multiple intrinsic ubiquitin receptors (i.e., Rpn1, Rpn10, or Rpn13). To test this, we integrated either wild type ubiquitin, ubiquitin^K48R^, or ubiquitin^K63R^ into the yeast genome at the URA3-Tim9 locus driven by the strong GPD promoter. The ubiquitin^K48R^ specifically reduces the length of K48-linked ubiquitin chains, the type of chain typically found on substrates targeted for proteasomal degradation (43). The ubiquitin^K63R^ specifically reduces the length of K63-linked ubiquitin chains, a modification that more commonly regulates signaling pathways or endocytosis of substrates (43). Crucially, reducing the length of K48-linked chains, but not K63-linked chains, interfered with the formation of proteasome condensates following azide treatment (Fig. 4B). Similarly, ubiquitin^K48R^ expression in the *ddi1*11 cells prevented the formation of proteasome condensates during logarithmic growth (Fig. 4C). A similar trend was observed for glucose starvation and prolonged growth (Fig. S3), however the manipulation of ubiquitin under those conditions might not be as effective (7) and manipulating the endogenous ubiquitin genes might be required to determine if K48-linked ubiquitin chains are involved under those conditions. Nevertheless, the notion that K48-linked ubiquitin chains are required for some proteasome condensates is supported by our observation that K48R-ubiquitin expression prevented proteasome condensates induction by azide as well as following antimycin A treatment, another inducer we previously characterized (Fig. S3) (39). In all, these data indicate that proteasome condensates in yeast consist of proteasomes, the shuttle factors Rad23 and Dsk2, and substrates with K48-linked chains.

### Different requirements for proteasome intrinsic receptors in condensate formation

Considering polyubiquitin chains serve as scaffolds holding proteasome condensates together, we would predict a critical role for the proteasome intrinsic ubiquitin receptors via binding to the ubiquitin chains and/or the shuttle factors’ UBL domains. However, a previous model proposed that proteasome storage granules consist of proteasome CP and RP in a dissociated state, together with the de-ubiquitinating proteasome associated factor Ubp6, and monoubiquitin (7). It is unclear in this model what forces induce condensate formation, but it presumably involves unidentified intrinsically disordered regions within some of the proteasome subunits and cellular pH as a potential trigger (7, 8). To distinguish between this and our model, we next analyzed the contribution of the proteasome ubiquitin receptors Rpn13, Rpn10, and Rpn1. To determine the role of the peripheral subunit Rpn13, we deleted the RPN13 gene, which does not impact overall proteasome stability or structure (44). We also deleted RPN10; however, this subunit plays a critical role in the stability of the RP, and its absence can result in some dissociation of the lid subcomplex. Therefore, we also introduced mutations, via CRISPR-Cas9 mutagenesis, that have previously been shown to strongly reduce the affinity of Rpn10 for ubiquitin, while not affecting proteasome structure (26). The Rpn1 subunit is essential, playing a structural role through its interaction with several other RP subunits. Therefore, we introduced a set of mutations that have been shown to disrupt ubiquitin binding (16). Analyzing these mutant strains, we observed surprising differences amongst the ubiquitin receptors for their role in proteasome condensate formation. The ability to efficiently form condensates following glucose starvation was clearly dependent on Rpn10, and perhaps modestly on Rpn13, while mutating Rpn1 had little to no impact on the ability of cells to form glucose starvation-induced condensates (Fig. 5A). However, when we analyzed azide-induced condensates, we noticed a strong dependance on Rpn13 and Rpn1 and little to no role for Rpn10 (Fig. 5B). All three ubiquitin receptors appear to play a role in the formation of condensates following prolonged growth in YPD (Fig. 5C). These data indicate a striking physiological and likely functional difference amongst these proteasome ubiquitin receptors. In all, our data show proteasome condensates are formed and maintained through a network of critical interactions amongst K48-linked ubiquitin chains, the shuttle factors Rad23 and Dsk2, and proteasomes (via the intrinsic proteasome ubiquitin receptors).

**Figure 5.**
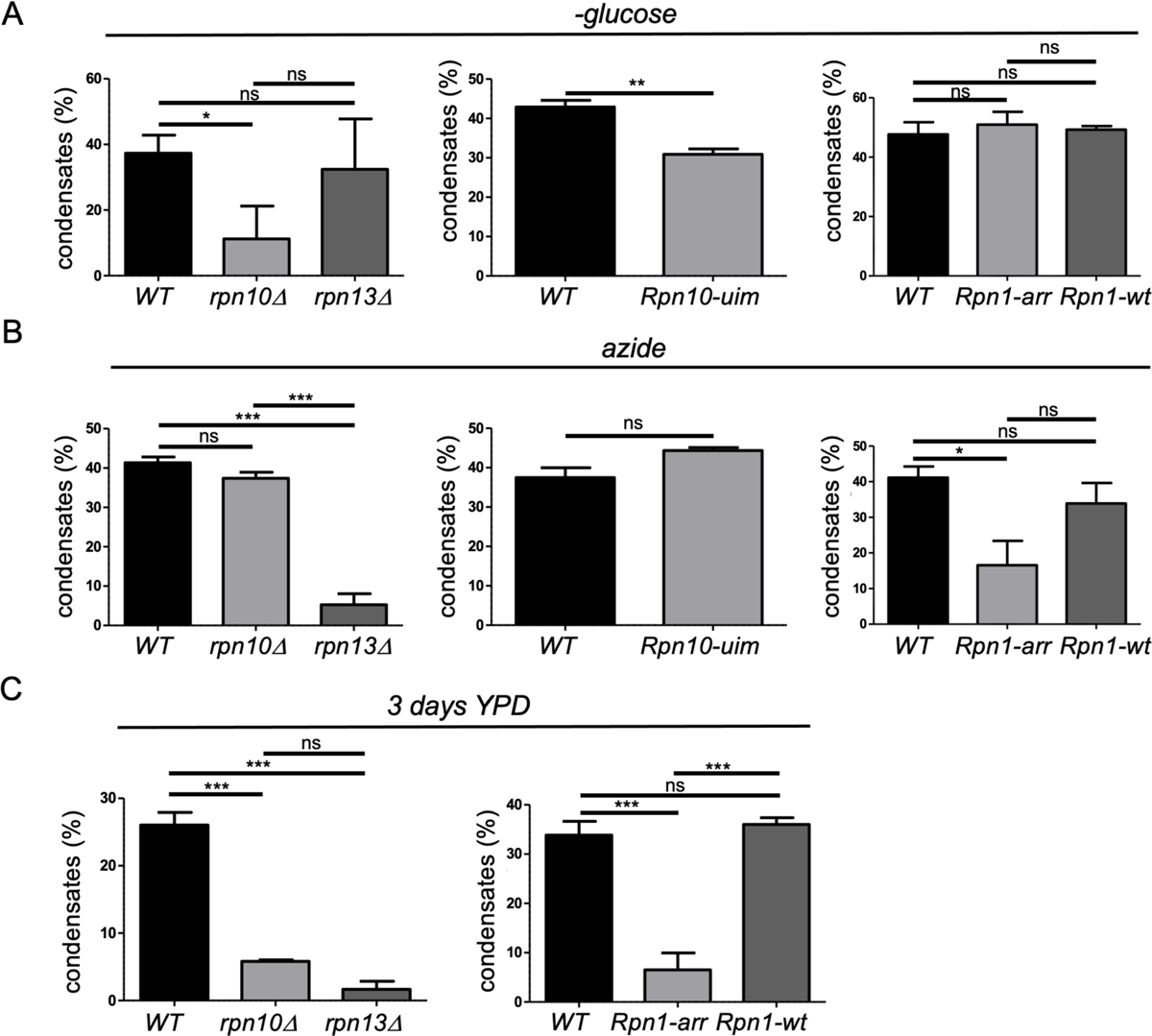
Intrinsic proteasome ubiquitin receptors have unique roles in proteasome condensate formation. **(A)** Left Panel, Rpn1-GFP expressing yeast deleted for Rpn10 or Rpn13 were starved for glucose and proteasome localization was monitored. The average of four independent experiments are presented with 1-way ANOVA and Tukey’s multiple comparison test used to determine significance. Center Panel, the ubiquitin interacting motif (UIM) of Rpn10 was mutated (Rpn10-uim). Cells were starved for glucose and proteasome localization monitored. Quantification from four independent experiments are presented with standard error of the mean. Paired t-test was used to determine significance. Right Panel, the ubiquitin interacting motif of Rpn1 was mutated (Rpn1-ARR) and compared to wildtype cells and cells with the same genetic modification, only lacking the ARR mutation (Rpn1-WT). Quantifications show the average of three independent experiments. 1-way ANOVA and Tukey’s multiple comparison test was used to determine significance. **(B)** The strains presented in A were treated with sodium azide to monitor the effects on proteasome condensates induced in this condition. First two panels show the results of four independent experiments with 1-way ANOVA and Tukey’s multiple comparison test used to determine significance for Rpn10 and Rpn13 mutants. SEM is presented for the Rpn10-uim mutant and T-test was used to determine significance. The right panel shows the results of 3 independent experiments and 1-way ANOVA and Tukey’s multiple comparison test was used to determine significance. **(C)** (left) Rpn1-GFP yeast deleted for RPN13 or RPN10 were grown for 3 days in YPD then imaged microscopically. 1-way ANOVA and Tukey’s multiple comparison test was used to determine significance. (right) Yeast as in A and B harboring WT Rpn1, Rpn1-arr, or Rpn1-wt were grown to stationary phase and monitored microscopically for condensate formation. 1-way ANOVA with Tukey’s multiple comparison test was used to determine significance. ns = not significant, * P < 0.05, ** P < 0.01, *** P < 0.001.

## Discussion

Proteasome condensates were first reported in yeast more than 15 years ago (described as PSGs). However, it has remained unclear what triggers their formation and what key interactions facilitate their liquid-liquid phase separation into condensates. Here, we report the presence of the proteasome shuttle factors Rad23 and Dsk2 in yeast proteasome condensates. We also found that long K48-linked ubiquitin chains are a key component of proteasome condensates in yeast, particularly upon inhibition of mitochondria and in *ddi1*11 cells (see summary Fig. 6). The disruption of condensate formation by the overexpression of ubiquitin^K48R^, which reduces the length of K48-linked ubiquitin chains, provides a strong indication these condensates contain long ubiquitin chains. Long chains would provide the capacity to serve as a scaffold, analogous to how mRNA works as a scaffold in stress granules, by allowing multiple binding partners to interact with the chain. The shuttle factors Rad23 and Dsk2 can interact with these chains utilizing their UBA domains. As the shuttle factors can bind to the proteasome intrinsic ubiquitin receptors (via their UBL domains) and ubiquitin chains, all components needed for multivalent interactions that allow condensate formation are present (Fig. 6). Indeed, mutating the intrinsic ubiquitin receptors of the proteasome disrupts proteasome condensate formation, indicating intrinsic receptors are critical for proteasome localization to these condensates. However, P62 and the human shuttle factors RAD23B and UBQLN2 have been shown to form condensates without proteasomes *in vitro* and/or *in vivo* (9, 38, 45, 46). For RAD23B K48-linked ubiquitin chains were more efficient then K63-linked in inducing LLPS *in vitro* (9). In contrast, for UBQLN2 condensate formation is more efficient with K63-linked ubiquitin chains as compared to K48-linked (47). However, neither of these *in vitro* studies included proteasomes in their assay. Shuttle factors could be the scaffolds or “stickers” that form the essence of the condensates. Proteasomes might thus be “clients” that enter in these condensates but are not critical for their formation. However, we don’t believe this to be the case because mutations of the intrinsic ubiquitin receptors do not cause an appearance of condensates that contain shuttle factors but lack proteasomes, instead it causes a general drop in condensates observed with either GFP-tagged proteasomes or mCherry-tagged Dsk2 (Fig. S4). Thus, the intrinsic receptors of the proteasome contribute critical multivalent interactions required for condensate formation.

**Figure 6.**
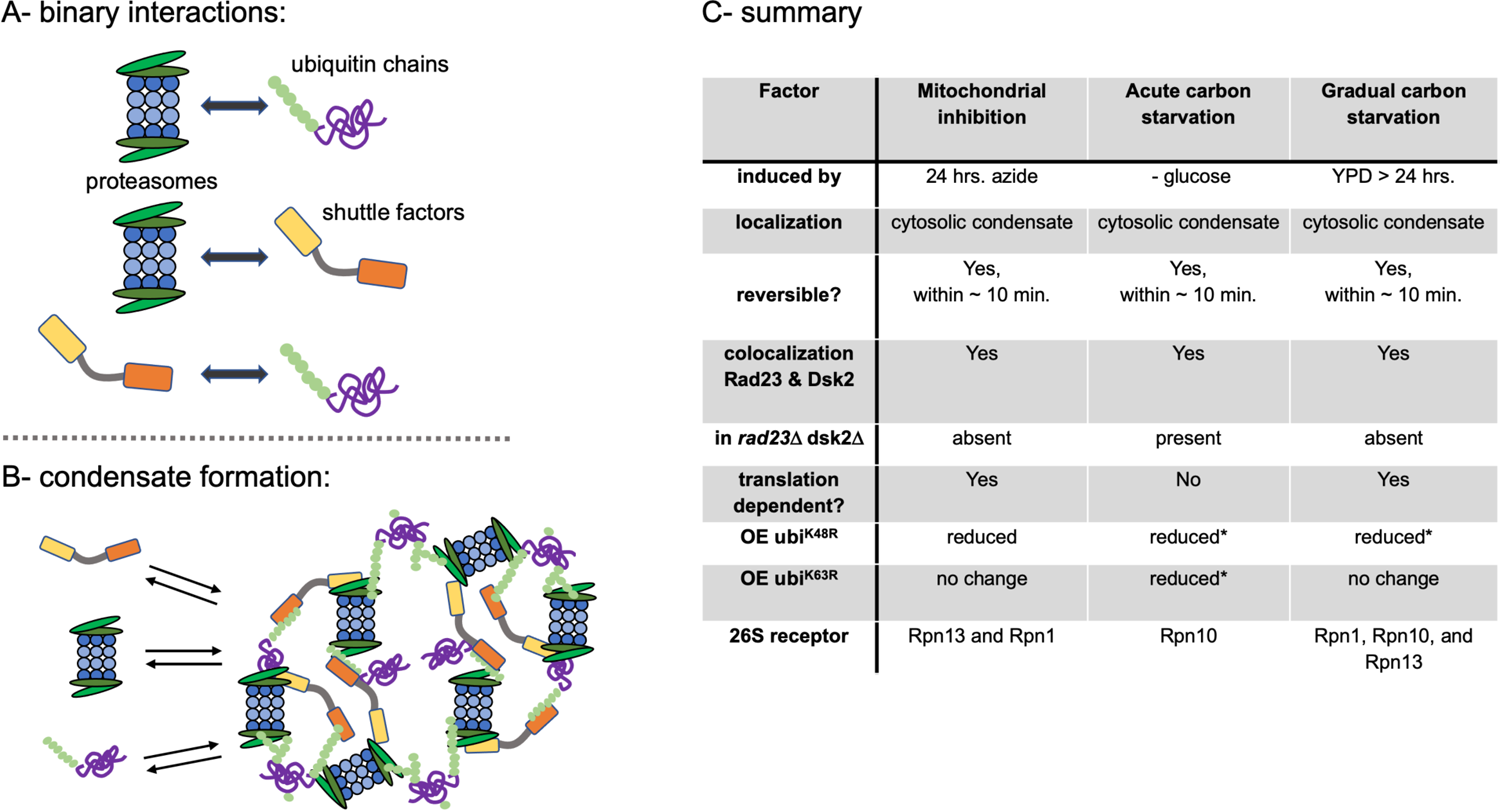
Model showing the multivalent interactions that drive proteasome condensate formation. **(A)** pairwise interactions that contribute to proteasome condensate formation. Proteasomes have three intrinsic ubiquitin receptors (Rpn1, Rpn10, Rpn13) that can interact with ubiquitin chains as well as Ubl domains of proteasome shuttle factors (yellow). Shuttle factors also have 1 or 2 UBA domains that can interact with ubiquitin chains (orange). (**B)** Our data show, at least for condensates induced prolonged growth or inhibition of mitochondria, that shuttle factors, K48-linked ubiquitin chains, and intrinsic ubiquitin receptors are all critical for condensate formation, suggesting that their concentration in combination with the network of interactions can drive condensate formation. **(C)** Summary of properties identified in this study for condensates formed under different stress conditions. Common features as well as distinguishing factors amongst these condensates are indicated. *\* shows trend, but is not statistically significant*.

Our data show that specific protein-protein interactions between these components are critical for condensate formation. This contrasts with a previous model that postulated an increase in monoubiquitin in cells triggers condensate formation in the cytosol, while polyubiquitinated proteins associate with and retain proteasomes in the nucleus (7). Our data suggest a more conserved general mechanism of the sequestration of proteasomes in condensates, as our data shows parallels to proteasome condensates in human cell (9-11). While the location and number of human proteasome condensates differ from yeast, their formation also involves a shuttle factor, RAD23B or p62, and polyubiquitin chains. In yeast, generally 1 or 2 larger condensates are observed in the cytosol, which is different from the many small condensates that have been observed in the nucleus of mammalian cells following specific stressors. Mammalian proteasome condensates have been proposed to be actively degrading substrates, suggesting there is an equilibrium between accumulating ubiquitinated substrates that enable LLPS versus the degradation of these substrates that counteracts this. In yeast, on the other hand, the condensates have been proposed to mature and store proteasomes, hence their name proteasome storage granules (PSGs) (6, 7, 48, 49). The difference in dynamics between human and yeast proteasome condensates could also explains why in general yeast cells end up with only one or two large, stable proteasome condensates while human cells have many small nuclear condensates. It remains unclear why proteasomes would need to be stored, as substrate selection and degradation are determined at the ubiquitination step. A proposed function of their storage has been to protect proteasomes from autophagy (8, 12). While attractive as a model, it should be noted that the majority of proteasomes are normally localized in the nucleus where they are already protected from autophagy (2, 4). Thus, it is unclear why proteasomes would be exported to the cytosol to protect them from autophagy. Either nuclear localization of proteasomes is unfavorable under those conditions, or the proteasome condensates have a function beyond protecting proteasomes. Interesting to note here is that we previously reported the involvement of a MAP kinase pathway in proteasome autophagy and the formation of proteasome condensates (3, 50). We proposed that this pathway was involved in regulating the nuclear export of proteasomes, suggesting that in addition to ubiquitinated substrates and shuttle factors, proteasomes need to be present at sufficient concentration in the cytosol to facilitate condensate formation.

The specific involvement of long K48-linked ubiquitin chains suggests that proteasome condensates include polyubiquitinated substrates. This is supported by our observation that a strain deleted for DDI1 shows condensates under conditions of logarithmic growth, i.e., without additional stress. In the absence of DDI1, cells accumulate substrates with long ubiquitin chains (41), suggesting that proteasome condensate formation can be triggered by the accumulation of substrates with long K48-linked ubiquitin chains. As the condensate-inducing conditions we tested in yeast all involve a drop in cellular energy levels (due to lack of carbon source or inhibition of mitochondrial oxidative phosphorylation), we hypothesize that such conditions lead to the accumulation of substrates in the cytosol, likely due to reduced proteolytic activity of the proteasome under those conditions, potentially combined with changes in activities of E3 ligases and deubiquitinating enzymes, something that remains to be explored. With the ubiquitin chain length and concentration as key triggers for condensate formation, we propose that the role of the condensates is to sequester ubiquitinated substrates, similar to how accumulated misfolded proteins are sequestered in membrane-less structures like IPODs or JUNQs. While these structures are less dynamic and contain aggregated material (49, 51, 52), many short-lived proteins targeted for degradation by ubiquitination are not misfolded but soluble and functional. Such proteins are unlikely to engage with the protein quality control machinery that would otherwise transport them to IPODs or JUNQs. Proteasome condensates could function to sequester these soluble ubiquitinated proteins and prevent them from remaining biologically active under conditions of low proteolytic activity. This would be similar to how stress granules sequester certain mRNAs to prevent translation under stress, thereby not spending energy producing superfluous proteins (53).

An interesting observation that remains to be further explored is why different condensate-triggering conditions required different proteasome intrinsic ubiquitin receptors: with condensates induced by glucose starvation depending on Rpn10 and those induced by azide depending on Rpn1 and Rpn13. These condensates also differ in their dependence on Rad23, Dsk2, and protein synthesis. Thus, there are clear distinguishing features between condensates induced by glucose starvation and those induced with mitochondrial stress. Interestingly, the degradation of substrates presented to the proteasome by shuttle factors or their UBL domain appears to be primarily mediated through Rpn1 and Rpn13 (54, 55). We hypothesize that different substrates accumulate based on the stressor, and those substrates interact differently with proteasome receptors. As the role of the different intrinsic ubiquitin receptors on the proteasome is still being debated, the dependance on different intrinsic receptors for specific conditions provides an exciting observation with significance beyond their role in proteasome condensate formation.

## Material and Methods

### Yeast strains and gene manipulations

Strains used in this work are reported in Table S1. Strains harboring different mutation or with C-terminal fluorescent fusions on proteasome subunits or shuttle factors were generated using standard PCR-based procedures (56, 57). Plasmids and primers used in this study are presented in table S2. All newly made plasmids were confirmed by sequencing. The DDI1 ORF was amplified from the yeast genome using primers pRL1106 and pRL1107 and cloned using NEBuilder® HIFI DNA assembly into pRS415 (CEN/LEU2), which had been amplified with pRL1108 and pRL1109, creating plasmid pJR980. Subsequent PCR based mutagenesis was performed to generate the protease dead D220N mutant plasmid (primers pRL1110 and pRL1111, plasmid pJR985). Yeast expressing these plasmids were grown overnight in selection media to maintain plasmid, then diluted in YPD for microscopic analysis after logarithmic growth. The URA-TIM9 genomic region was used to integrate wildtype and mutant ubiquitin. First, Ubiquitin driven by the strong GDP promoter and followed by a ADH1 terminator and HIS3 MX6 selection cassette was cloned into a pNC1124 (addgene #41560, (58)) derived plasmid such that it was flanked by region from the URA3 and Tim9 genes that serve for cross over. The resulting plasmid, pJR1040, was mutated to create Ubiquitin^K48R^ (pJR1048) or Ubiquitin^K63R^ (pJR1050). For integration of this cassette into strains backgrounds with a mutated URA3, the plasmids were digested with SalI and SacI to create a linear fragment that was utilized in yeast transformation. For backgrounds with a clean deletion of the URA3 gene (such as the yeast knock-out collection), which lack the URA3 flanking region for cross over, PCR with primers pRL995 and pRL559 was used to create a linear fragment for transformation. CRISPR-Cas9 mutagenesis was used to generate yeast harboring mutation in the Rpn10 ubiquitin interacting motif (rpn10-uim, replacing LAMAL with NNNNN). Sequence coding for the guide RNA (GAACTGGCAATGGCCTTGCG) was cloned into CRISPR vector pML107 (Addgene plasmid # 67639 (59)), creating plasmid pJR979. pJR979 and the repair duplex (see table S1) were transformed into yeast and mutations were confirmed by sequencing the genomic region. To introduce the mutation for Rpn1 in the genomic RPN1 locus, we digested plasmid YSp97 (for WT control) or pEL356c (encoding *RPN1-ARR* mutation) with XhoI and NotI (16). Linear fragments were transformed into yeast and successful integration of the mutant RPN1 was confirmed by sequencing the genomic region.

### Yeast Growth Conditions

Yeast strains were inoculated in yeast extract peptone media containing dextrose as a carbon source (YPD) and grown overnight. For starvation, sodium azide treatment or antimycin A treatment, cultures were diluted to OD600 = 0.5 in fresh media and grown for 4 hours. Sodium azide was added to a final concentration of 0.5 mM and antimycin A at a concentration of 0.1 mM. For starvation, logarithmically growing cultures were harvested, washed, and inoculated in either SD media lacking carbon or nitrogen as previously described (60). The proteasome inhibitor PS-341 (also known as Bortezomib or Velcade) was added at a final concentration 100 μM. For cycloheximide treatment, cells were first pre-treated with cycloheximide for 20 min. before inoculation in final growth condition: overnight cultures were diluted in fresh YPD as above, grown for four hours, and cycloheximide was added to a final concentration of 50 µg/mL. Cultures were incubated for 20 min, then cells were washed with respective media, inoculated at an OD600 of 1.5 in respective media containing 50 µg/mL cycloheximide, and allowed to grow to indicated time points.

### Fluorescence microscopy

Live yeast cells were collected by centrifugation (2,000 g, 2 min.) and resuspended in PBS buffer, or a small volume of supernatant. Cells were immobilized on a microscopy slide using a 1% agarose pad supplemented with PBS buffer (modified from https://www.youtube.com/watch?v=ZrZVbFg9NE8, 2019). Images were acquired at room temperature on a Nikon Eclipse TE2000-S microscope at ×600 magnification using a Plan Apo ×60/1.40 objective and R3 Retiga camera. For GFP images, Sedat Quad filter set (Chroma 86000v2, Bellows Falls, VT) was set to an excitation wavelength of 490/20 nm and emission wavelength of 528/38 nm. mCherry images were obtained using excitation and emission wavelengths of 555/28 nm and 685/40 nm, respectively. Images were collected using Metamorph (Molecular Devices) within 10 minutes of cell immobilization on microscopy slides. Fluorescent signal was analyzed and measured using FIJI. For proteasome granule quantification, over 100 cells per experimental replicate were counted and scored (granules/no granules). Scale bars represent 5 μM.

### Cell lysis & immunoblotting analysis

At indicated time points, for each strain, an A600 of 2 was harvested by centrifugation and immediately lysed or stored at −80 °C. Alkaline lysis was carried out as previously reported (61). In short, pellets were resuspended in 100 μL distilled water to which 100 μL 200 mM NaOH was added, and samples were incubated at room temperature for 5 min. Cell suspensions were pelleted, resuspended in 50 μL SDS-PAGE sample buffer (0.06 M Tris–HCl, pH 6.8, 5% glycerol, 2% SDS, 4% β-mercaptoethanol, 0.0025% bromophenol blue), boiled at 96 °C for 5 min, and supernatant was collected. SDS-PAGE of these samples was followed by Western blotting. Membranes were analyzed by immunoblotting using antibodies against GFP (Roche Applied Science, catalog no.11814460001), Pgk1 (Invitrogen, catalog no.459250) and ubiquitin (Vu1, LifeSensors, catalog no. VU101). Images were acquired using a Gbox imaging system (Syngene) and captured with GeneSnap software.

## Acknowledgement

We want to thank members of the Roelofs lab and Drs. Jeremy Schmidt, Daniel Kraut, and Carlos Castañeda for helpful discussions and feedback on the manuscript. This work was supported by a grant from NIH-NIGMS (R01GM118660 and K-INBRE grant P20GM103418).

## Author contributions

All authors designed research; K.A.W., G.V., and S.Y.L. performed research and analyzed data and made figures; J.R. wrote the paper with input from all authors.

## Lead Contact and Materials Availability

Further information and requests for resources and reagents should be directed to and will be fulfilled by the Lead Contact, Jeroen Roelofs.

## Data, Materials, and Software Availability

All study data are included in the article and/or SI Appendix.

## Competing interests

The authors declare no competing interest.

## Notes

### Competing Interest Statement

The authors have declared no competing interest.

