## Supplemental figures and tables for "Proteasome condensate formation is driven by multivalent interactions with shuttle factors and K48-linked ubiquitin chains"

### for

[2] current address: LSUHSC, Department of Genetics; Louisiana Cancer Research Center

[*] equal contributions

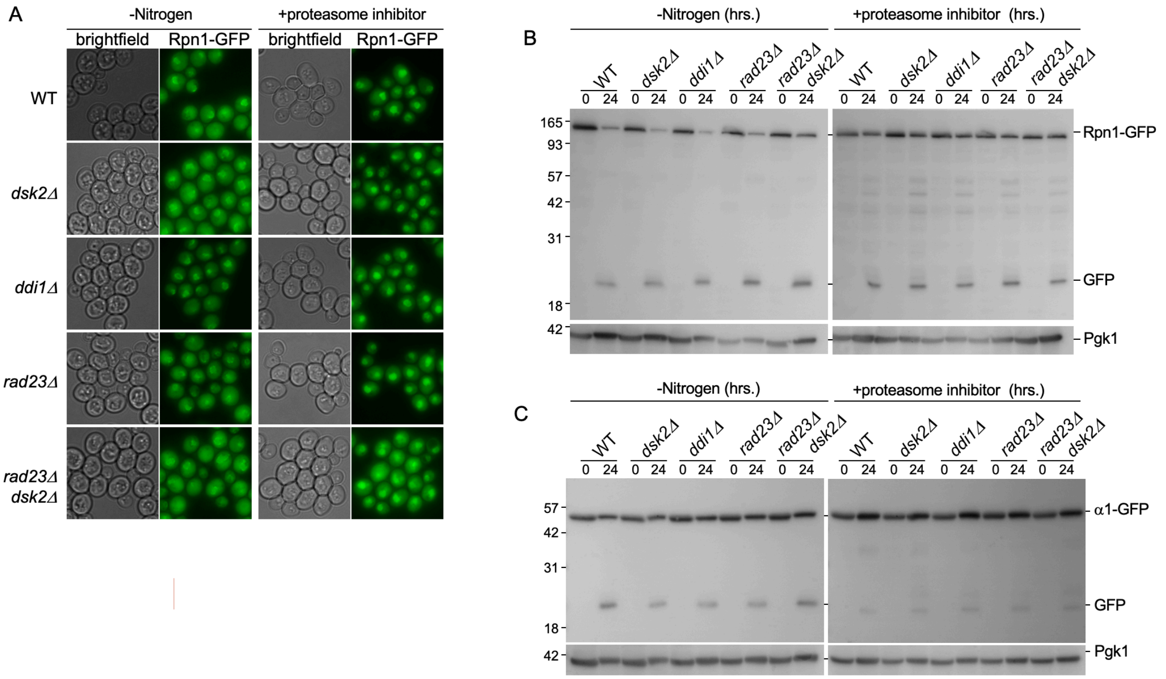


**Figure S1. Yeast shuttle factors are not required for proteasome autophagy.**

(A) Wild type, *dsk2*Δ, *ddi1*Δ, *rad23*Δ and *rad23*Δ *dsk2*Δ yeast expressing Rpn1-GFP were starved of nitrogen or treated with proteasome inhibitor for 24 hours to monitor proteasome autophagy. (B) Western blots to monitor the accumulation of "free GFP" as a readout for proteasome autophagy. Samples were collected from cells as in (A) and lysates blotted for GFP and loading control Pgk1. (C) Wild type and shuttle factor mutants expressing α1-GFP were used to monitor autophagy of the proteasome core particle.

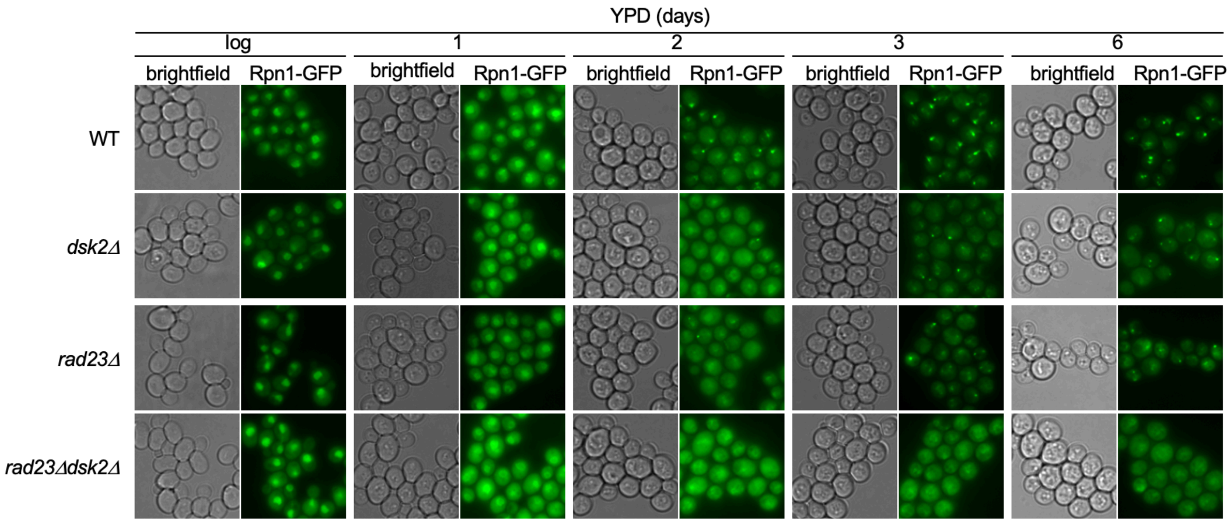


**Figure S2. Rad23 and Dsk2 are required for proteasome condensate formation in quiescence.** Wild type, *dsk2*Δ, *rad23*Δ and *rad23*Δ *dsk2*Δ yeast expressing Rpn1-GFP were grown for the indicated times in YPD media then monitored microscopically for proteasome condensate formation.

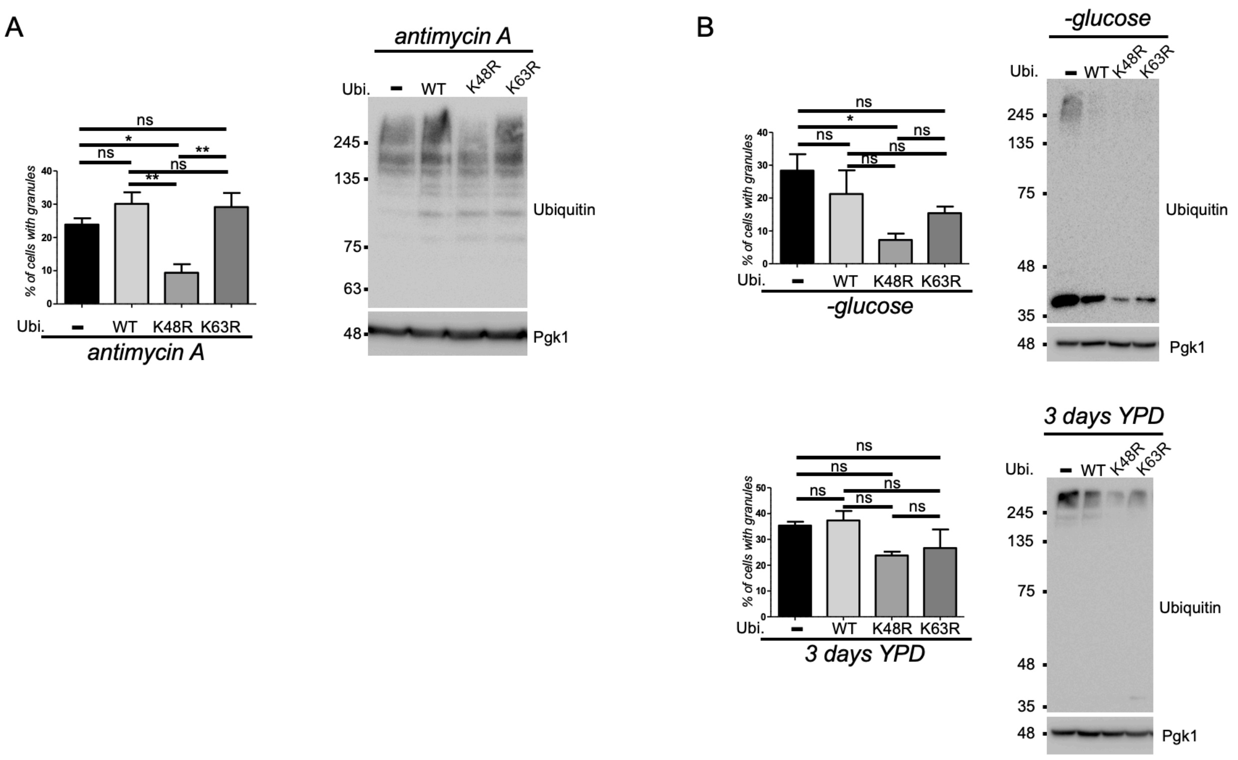


**Figure S3. The dependance of proteasome condensate on long K48-linked and K63-linked ubiquitin chains under different condensate inducing conditions.**

**(A)** Rpn1-mCherry yeast expressing no additional ubiquitin, WT ubiquitin, K48R ubiquitin or K63R ubiquitin were treated with antimycin A for 24 hrs and analyzed for proteasome condensate formation (left panel). The average of four independent experiments is shown and 1-way ANOVA with Tukey's multiple comparison test was used to determine significance. Right panel shows immunoblots of total lysate for ubiquitin and loading control Pgk1. **(B)** Same analysis following 24 hrs. of glucose starvation or prolonged growth in YPD. ns = not significant, * $P < 0.05$.

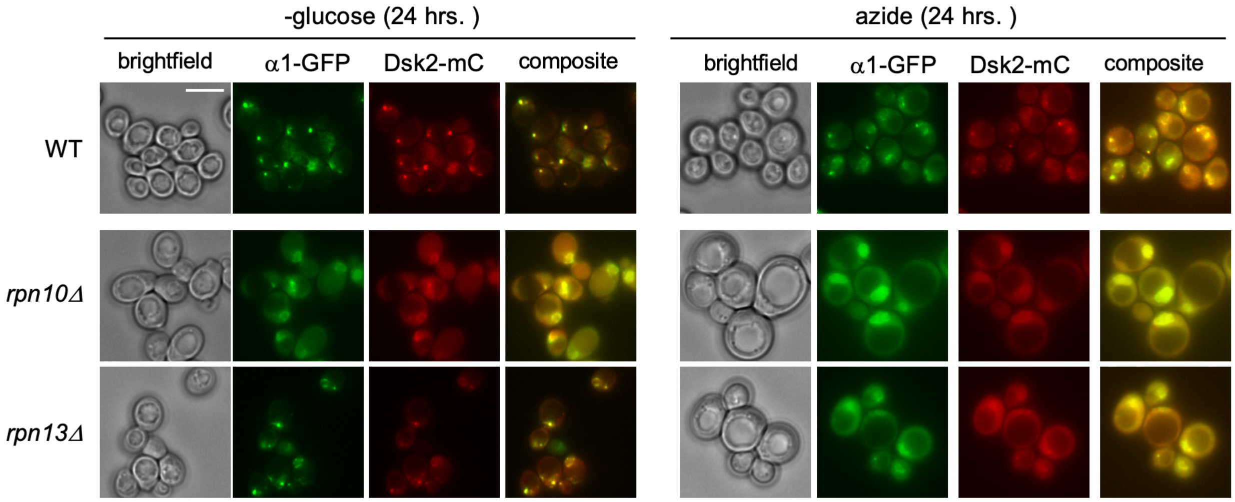


**Figure S4. Mutation of proteasome intrinsic receptors disrupts condensate formation for proteasomes and the shuttle factor Dsk2.**

α1-GFP, Dsk2-mCherry expressing yeast deleted for Rpn10 or Rpn13 were starved for glucose or treated with sodium azide for 24 hours. Microscopic analysis shows that when proteasomes fail to form condensates, Dsk2 also fails to condensate. The scale bar represents 5 μm.

#### **Supplementary Table 1. Primer list.**

| Primer | Genotype | Template | Sequence (5' to 3') |
|---|---|---|---|
| Rvrs/Rpn1 | *rpn1::rpn1-GFP(-HIS)/rpn1-GFP(CloNat)* | pYM28/pJR788 | TTTGAATTTTTCCTATTCTGGTTGATATTGCCCAAAAGCTATTCAGTTTAATCGATGAATTCGAGCTCG |
| Frwd/Rpn1 | *rpn1::rpn1-GFP(-HIS)/rpn1-GFP(CloNat)* | pYM28/pJR788 | TTGAGGGCGTAGTAATTTTAAAGAAGAACCCTGACTATCGTGAAGAGGAGCGTACGCTGCAGGTCGAC |
| pRL608 | *Dsk2-mCherry* | pBSS35 | CGTCCAAGGCGCTCTTGATTCACTACTGAACGGCGATGTTGGTCGACGGATCCCCGGG |
| pRL609 | *DSK::G418* | pFA6a-kanMX6 | GAGTAGGGTAAAAGTATATAGGTTGCGGCATCTAGACGTTATCGATGAATTCGAGCTCG |
| pRL612 | *Rad23-mCherry* | pBSS35 | AGAAGCTGCAGCAAATATTCTATTCAGCGATCATGCCGACGGTCGACGGATCCCCGGG |
| pRL616 | *Ddi1-mCherry* | pBSS35 | CAGACTAACGGAAATGCAGAATTTGCTGCATCCCTCCTTTTCCAAGGTCGACGGATCCCCGGG |
| pRL621 | *DSK::G418* | pFA6a-kanMX6 | GCAAATAAGACGGATCAAAGACACCGAATCATTCTAGCACGCGTACGCTGCAGGTCGACG |
| pRL613 | *RAD23::G418* | pFA6a-kanMX6 | GTCGAGAATAAGTGAAGATACTTCAAGCCATAACATTACTATCGATGAATTCGAGCTCG |
| pRL620 | *RAD23::G418* | pFA6a-kanMX6 | CTTTAAATCACAGATCACACAAAGACAACATACAATAGAACGTACGCTGCAGGTCGACG |
| pRL617 | *DDI1::G418* | pFA6a-kanMX6 | GGGCTACATACGTAGAGGCCGATCACAATATCAGTGGTTGATCGATGAATTCGAGCTCG |
| pRL622 | *DDI1::G418* | pFA6a-kanMX6 | AAAGTACATACCAAACATAACAGCAAAAATATACGTAAAGCGTACGCTGCAGGTCGACG |
| pRL36 | *SCL1::SCL1-mCherry* | pBSS35 | GTGTTGACGCGTGTGATTTCACATTATGTTGTGGCAGGAAGATCGATGAATTCGAGCTCG |
| pRL710 | *SCL1::SCL1-mCherry* | pBSS35 | TGCTGAGAACATCGAAGAAAGGCTAGTAGCAATTGCTGAACAAGATGGTCGACGGATCCCCGGG |
| pRL1154 | *Rpn13::HYGRO* | pFA6a-hph | CCTAAGTGCTGGTTGACTTATAAATTTTAAAAGAGTGTTG CGTACGCTGCAGGTCGAC |
| pRL1155 | *Rpn13::HYGRO* | pFA6a-hph | TTCTCTTCAGTTTTTTATCAAAAAATGCCACAACCTTGAT ATCGATGAATTCGAGCTCG |
| pRL1073 | repair duplex for *Rpn10-UIM* | | 5'-TGGACTTCGGGGTAGACCCATCAATGGACCCAGAAAACAACAATAATAACCGTCTGTCTATGGAAGAAGAGCAGCAAAGACAGGAAAG-3'<br>3'-ACCTGAAGCCCCATCTGGGTAGTTACCTGGGTCTTTTGTTGTTATTATTGGCAGACAGATACCTTCTTCTCGTCGTTTCTGTCCTTTC-5' |
| pRL1106 | *DDI1* gene | *Sc* gDNA | CTTGAGTCTCCTTCAAGTACAACG |
| pRL1107 | *DDI1* gene | *Sc* gDNA | GTATGGGGTTGCATGTCATTCGG |
| pRL1110 | *Ddi1$^{D220N}$* | pJR980 | AATACAGGGGCTCAAACAACG |
| pRL1111 | *Ddi1$^{D220N}$* | pJR980 | TACAAATGCCTTTACGGGGTAG |


1. Goldstein, A. L. & McCusker, J. H. Three new dominant drug resistance cassettes for gene disruption in Saccharomyces cerevisiae. Yeast 15, 1541–53 (1999).
2. Janke, C. et al. A versatile toolbox for PCR-based tagging of yeast genes: new fluorescent proteins, more markers and promoter substitution cassettes. Yeast 21, 947–62 (2004).
3. Hailey DW, Davis TN, Muller EG. Fluorescence resonance energy transfer using color variants of green fluorescent protein. Methods Enzymol. 351:34-49 (2002.

#### Supplementary Table 2. Strain list.

| Strain | Genes manipulated | Figure(s) | Ref. |
|---|---|---|---|
| sJR1255 | *RPN1::RPN1-GFP* (HIS3) | Fig.1B,C,D,E; Fig.3A,D; S1A,B;S2 | b |
| sJR1123 | *RPN1::RPN1-GFP* (HIS3) *DSK2::KAN* | Fig.1B,C,D; S1A,B; S2 | b |
| sJR1124 | *RPN1::RPN1-GFP* (HIS3) *DDI1::KAN* | Fig.3A,B,D; Fig. 4A; S1A,B | b |
| sJR1127 | *RPN1::RPN1-GFP* (HIS3) *RAD23::KAN* | Fig.1B,C,D; S1A,B; S2 | b |
| sJR1143 | *RPN1::RPN1-mCherry* (G418) | Fig. 4B; S3 | b |
| sJR1203 | *RPN1::RPN1-GFP* (HIS3) *RAD23::G418 DSK2Δ::cloNAT* | Fig.1B,C,D; S1A,B;S2 | b |
| sJR861 | *RPN1::RPN1-GFP* (HIS3) | Fig. 5A,B, C | (2) |
| sJR897 | *RPN1::RPN1-GFP* (HIS3) *RPN10*::HYGRO | Fig. 5A,B, C | (2) |
| sJR1084 | *SCL1::SCL1-GFP* (HIS3) | Fig. 5A,B, S4 | (3) |
| sJR1256 | *SCL1::SCL1-GFP* (HIS3) | Fig.1B,C,D; S1A,C | b |
| sJR1128 | *SCL1::SCL1-GFP* (HIS3) *RAD23::G418* | Fig.1B,C,D; S1A,C | b |
| sJR1137 | *SCL1::SCL1-GFP* (HIS3) *DSK2::G418* | Fig.1B,C,D; S1A,C | b |
| sJR1138 | *SCL1::SCL1-GFP* (HIS3) *DDI1::G418* | Fig.3A, S1A,C | b |
| sJR1284 | *SCL1::SCL1-GFP* (HIS3) *RAD23::G418 DSK2Δ::cloNAT* | Fig.1B,C,D; S1A,C | b |
| sJR1263 | *RPN1::RPN1-GFP* (HIS3) *RAD23::KAN DSK2Δ::cloNAT DDI1Δ::HYG* | Fig.3D | b |
| sJR1294 | *RPN1::RPN1-GFP* (HIS3) *RAD23::KAN DDI1Δ::HYG* | Fig.3D | b |
| sJR1299 | *RPN1::RPN1-GFP* (HIS3) *DSK2Δ::cloNAT DDI1Δ::HYG* | Fig.3D | b |
| sJR1052[a] | *RPN1::RPN1-GFP* (cloNAT) | Fig. 4A | b |
| sJR1208[a] | *RPN1::RPN1-GFP* (cloNAT) DSK2::DSK2-mCherry (HYGO) | Fig. 2 | b |
| sJR1210[a] | *RPN1::RPN1-GFP* (cloNAT) DDI1::DDI1-mCherry (HYGO) | Fig. 3C | b |
| sJR1222[a] | *RPN1::RPN1-GFP* (cloNAT) RAD23::RAD23-mCherry (HYGO) | Fig. 2 | b |
| sJR1903 | *RPN1::RPN1-GFP* (cloNAT) pJR980 (DDI1-WT) | Fig. 4A | b |
| sJR1904 | *RPN1::RPN1-GFP* (cloNAT) pJR985 (DDI1-D220N) | Fig. 4A | b |
| sJR1905 | *RPN1::RPN1-GFP* (HIS3) *DDI1::KAN* pJR980 (DDI1-WT) | Fig. 4A | b |
| sJR1906 | *RPN1::RPN1-GFP* (HIS3) *DDI1::KAN* pJR985 (DDI1-D220N) | Fig. 4A | b |
| sJR2112 | *RPN1::RPN1-mCherry* (G418) GPD-WT Ubiquitin (HIS) | Fig. 4B; S3 | b |
| sJR2113 | *RPN1::RPN1-mCherry* (G418) GPD-K48R Ubiquitin (HIS) | Fig. 4B; S3 | b |
| sJR2114 | *RPN1::RPN1-mCherry* (G418) GPD-K63R Ubiquitin (HIS) | Fig. 4B; S3 | b |
| sJR2227 | *RPN1::RPN1-GFP* (HIS3) *RPN13::HYGRO* | Fig. 5A,B, C | b |
| sJR2247 | *RPN1::RPN1-GFP* (HIS3) *RPN10::Rpn10-uim* | Fig. 5A,B | b |
| sJR2147 | *RPN1::RPN1-GFP* (HYGRO) GPD-WT Ubiquitin (HIS) | Fig. 4C | b |
| sJR2149 | *RPN1::RPN1-GFP* (HYGRO) *DDI1::G418* GPD-WT Ubiquitin (HIS) | Fig. 4C | b |
| sJR2152 | *RPN1::RPN1-GFP* (HYGRO) GPD-K48R Ubiquitin (HIS) | Fig. 4C | b |
| sJR2154 | *RPN1::RPN1-GFP* (HYGRO) *DDI1::G418* GPD-K48R Ubiquitin (HIS) | Fig. 4C | b |
| sJR2253 | *SCL1::SCL1-GFP* (HIS3) *RPN1::RPN1-ARR* (TRP1) | Fig. 5A,B, C | b |
| sJR2254 | *SCL1::SCL1-GFP* (HIS3) *RPN1::RPN1-WT* (TRP1) | Fig. 5A,B, C | b |
| sJR2261 | *SCL1::SCL1-GFP* (HIS3) DSK2::DSK2-mCherry (G418) | S4 | b |
| sJR2262 | *SCL1::SCL1-GFP* (HIS3) DSK2::DSK2-mCherry (G418) *RPN10*::HYGRO | S4 | b |
| sJR2263 | *SCL1::SCL1-GFP* (HIS3) DSK2::DSK2-mCherry (G418) *RPN13::HYGRO* | S4 | b |

*genetic background is MATα his3Δ1 leu2Δ0 lysΔ0 ura2Δ0 (BY4172).*
*(a) genetic background: MATa can1Δ::STE2pr-Sp_his5 lyp1Δ his3Δ1 leu2Δ0 ura3Δ0 met15Δ0*

*(b)) this study.*
